# Multiplexed live-cell visualization of endogenous proteins with nanometer precision by fluorobodies

**DOI:** 10.1101/145698

**Authors:** Alina Klein, Susanne Hank, Anika Raulf, Felicitas Tissen, Mike Heilemann, Ralph Wieneke, Robert Tampé

## Abstract

The visualization of endogenous proteins in living cells is a major challenge. A fundamental requirement for spatiotemporally precise imaging is a minimal disturbance of protein function at high signal-to-background ratio. Current approaches for visualization of native proteins in living cells are limited by dark emitting, bulky fluorescent proteins and uncontrollable expression levels. Here, we demonstrate the labeling of endogenous proteins using nanobodies with site-specifically engineered bright organic fluorophores, named fluorobodies. Their fast and fine-tuned intracellular transfer by microfluidic cell squeezing allowed for low background, low toxicity, and high-throughput. Multiplexed imaging of distinct cellular structures was facilitated by specific protein targeting, culminating in live-cell super-resolution imaging of protein networks. The high-throughput delivery of engineered nanobodies will open new avenues in visualizing native cellular structures with unprecedented accuracy in cell-based screens.

## Introduction

The localization of endogenous proteins in native settings is fundamental for quantitative spatiotemporal understanding of cellular homeostasis and dysfunction. Here, live-cell imaging is highly desired to visualize timely-balanced protein networks, avoiding contingent artifacts by fixation or permeabilization procedures.^1^ In this regard, genetic fusion of fluorescent proteins (FPs) enabled the visualization of proteins of interest (POIs).^2^ However, the fusion to FPs or other bulky enzyme-based tags (*e.g.* SNAP_f_ tag)^3^ can provoke protein misassembly, mistargeting, and overexpression artifacts.^4^ Alternatively, antibodies and recombinant binders are prominent tools to trace proteins.^5-8^ Yet, targeting with conventional antibodies is limited to fixed cells since their disulfide bonds are reduced in the cytosol. To overcome this downside, antibody fragments have been developed for protein targeting.^9,10^ For in-cell labeling of endogenous proteins, the binder must fulfill several demands: high affinity and specificity to its target, long-term stability in the reducing milieu of the cytosol, and small size avoiding interference with the POI’s function. In this light, the variable domains of heavy-chain only antibodies (V_H_Hs) of camelids, named nanobodies, are prime candidates.^11,12^ They bind with nanomolar or even picomolar affinities to their targets, are very small in size (~ 13 kDa), exhibit great solubility, and can be functionally produced in the reducing cytosolic milieu of mammalian cells. Nanobodies fused to fluorescent proteins (chromobodies) have been used to trace various POIs.^12-18^ Recently, a chromobody has been developed for actin labeling in the cytosol and nucleus of mammalian cells,^19^ surpassing common techniques such as LifeAct by minimizing interference with actin dynamics.^20^ Nonetheless, visualization of intracellular proteins is still impeded because fused FPs triple the size of the nanobody and FPs entail suboptimal photophysical properties for super-resolution microscopy. An additional drawback of chromobodies is that their expression level cannot be precisely controlled, evoking high background and deteriorated signal-to-background staining, which in turn makes the observation of low abundant proteins and in particular super-resolution microscopy very difficult or impossible. As recently demonstrated, nanobodies directed against FPs were equipped with quantum dots or organic dyes to trace kinesin by single-particle tracking.^21,22^ However, both studies were limited by the expression of a genetically modified POI.

So far, methods visualizing endogenous proteins in live unmodified cells have not been accomplished. To overcome this limitation, we advanced nanobodies with bright, high-performance fluorescent probes, named fluorobodies. Combined with high-throughput cellular delivery by a vector-free microfluidic device,^23,24^ endogenous proteins were visualized by multi-color imaging at excellent signal-to-background ratios. Ultimately, the endogenous nuclear lamina was resolved at nanometer precision by live-cell super-resolution microscopy.

## Results

### Specific protein labeling *via* fluorobodies

As a first showcase, we modified the established α-GFP nanobody^25,26^ by strategic positioning of small organic fluorescent probes. We engineered nanobodies with a single free cysteine either at the N or C terminus, or position 9, respectively. After expression and affinity purification, the nanobodies were labeled via maleimide coupling with high-quantum yield fluorophores of, *e.g.* sulfo-Cyanine 3 (sCy3), sCy5, or ATTO655 (**Figure 1a**). To suppress transient exposure of inner scaffold cysteines by thermal breathing, covalent dye attachment was performed at 4 °C.^27^ Specific conjugation of the fluorophore was confirmed by SDS-PAGE in-gel fluorescence and fluorescence-detection size-exclusion chromatography, with molar labeling ratios of ~ 0.9 for α-GFP^sCy3^ and ~ 0.8 for α-GFP^sCy5^ (**Figure 1b** and **Figure 1 – Figure Supplement 1**).

**Figure 1.**
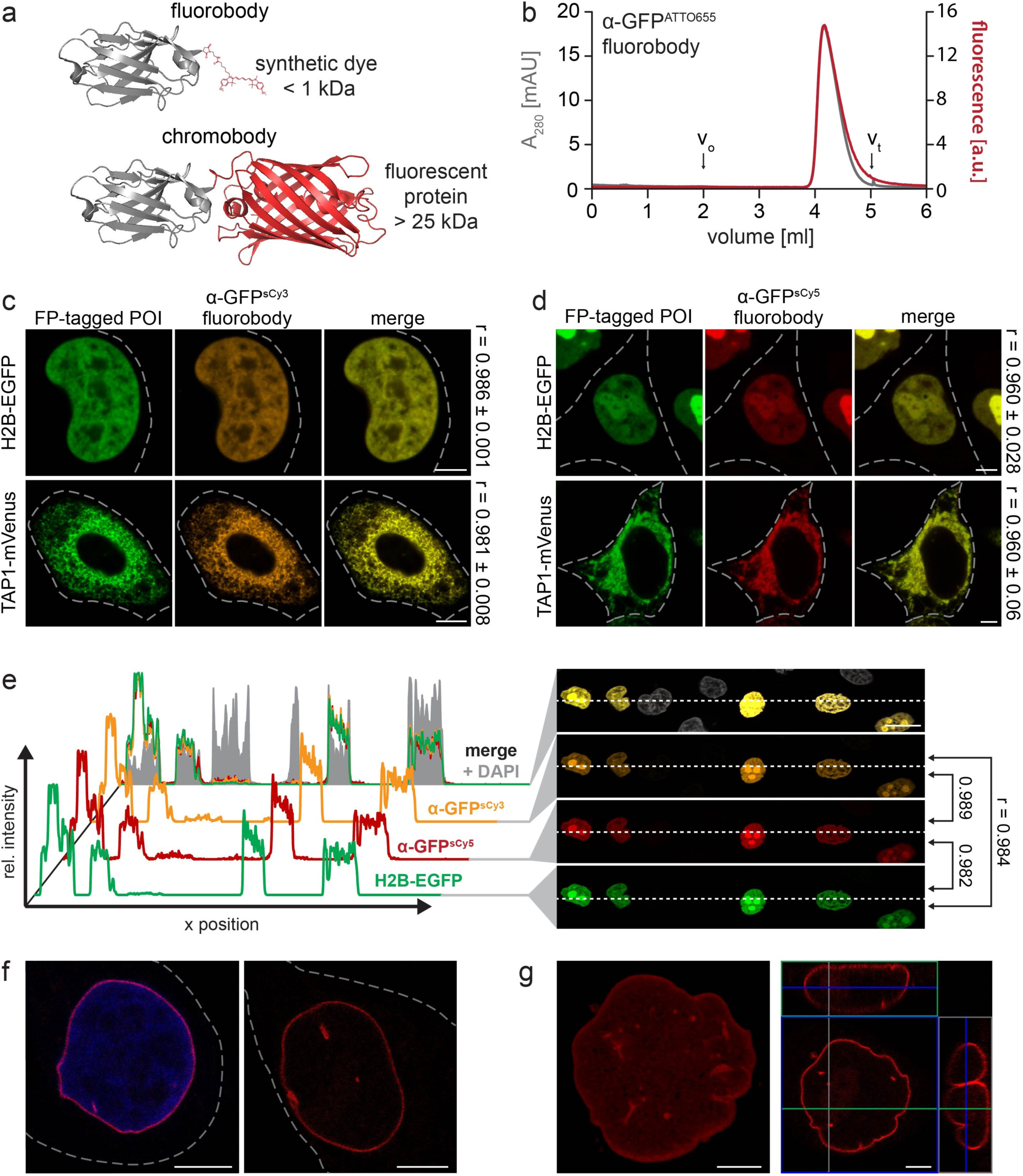
Ι Specific binding of site-specifically labeled fluorobodies. (**a**) Size comparison of nanobodies labeled with a fluorescent protein (chromobody) or a synthetic dye (fluorobody). Bulky fluorescent proteins (> 25 kDa) triple the size of the nanobody (~13 kDa). In comparison, the minimal size of fluorophores (< 1 kDa) is unlikely to disturb the nanobody and offer a higher quantum yield as well as photostability. mCherry (PDB 2H5Q) is shown exemplarily. (**b**) Size-exclusion chromatography (SEC) of the site-specifically labeled α-GFP^ATTO655^ Fb. The grey and red lines represent absorption at 280 nm (A_280_) and ATTO655 fluorescence at λ_ex/em_ 655/680 nm. Chromatogram shows specific modification of the nanobody and high purity. (**c, d**) Specific labeling of FP-tagged proteins in chemically arrested cells. HeLa Kyoto cells expressing H2B-EGFP or TAP1-mVenus (green) were fixed with 4% formaldehyde and labeled with 100 nM of α-GFP^sCy3^ (**c**, orange) or α-GFP^sCy5^ (**d**, red) Fb. Excellent co-localization (merge) between the Fbs and both FP-tagged POIs was detected. Pearson’s coefficients (r) were calculated from 9-11 individual cells (right). (**e**) Fixed HeLa Kyoto cells expressing H2B-EGFP were stained with α-GFP^sCy3^ (orange), α-GFP^sCy5^ (red, 100 nM each), and DAPI (blue). Cross-sections of relative fluorescence intensity profiles of both Fbs (along dashed line) highly correlated with the expression level of H2B-EGFP. Low background fluorescence was detected in untransfected cells (only DAPI positive), supported by the Pearson’s coefficients (r) between different channels (right). (**f, g**) After fixation, the endogenous nuclear lamina of HeLa Kyoto cells was visualized by 100 nM of α-Lamin^ATTO655^ Fb (red). (**g**) The Fb specifically decorated the nuclear envelope (DNA optionally visualized via DAPI staining, blue). (**f**) 3D reconstruction of a z-stack showed a dense labeling of the nuclear lamina along with a high signal-to-background ratio (see Supplementary Video 1). Images were taken by CLSM with the Airy Scan detector (f and g). Dashed lines indicate the cell border. Scale bars: 5 µm (**c**,**d,f,g**) and 20 µm (**e**).

As an initial screen, we evaluated specific binding of the α-GFP fluorobodies (Fbs) in fixed cells prior to live-cell applications. Different intracellular assemblies were selected as targets, *e.g.* the transporter associated with antigen processing TAP (TAP1-mVenus) in the ER membrane, histone 2B (H2B-EGFP) in the nucleus, and lamin A (mEGFP-Lamin A) as a component of the nuclear envelope. The respective POIs were expressed in human HeLa Kyoto cells, labeled with the α-GFP Fbs (50-200 nM) and analyzed by confocal laser scanning microscopy (CLSM). The Fbs retained high target specificity with negligible background staining (**Figure 1c,d** and **Figure 1 – Figure Supplement 2**). More important, an excellent co-localization was observed as depicted by Pearson’s coefficients ranging from 0.948 to 0.986, independently of the attached synthetic dyes (sCy3 and sCy5; **Figure 1c,d** and **Figure 1 – Figure Supplement 2**). The Fb intensity distribution highly correlated with the POI expression level, reflecting specific and stoichiometric binding (**Figure 1e** and **Figure 1 – Figure Supplement 2** and **3**).

To avoid limitations typically entailed by overexpression of fusion proteins, we traced the endogenous nuclear envelope by an engineered α-Lamin Fb. As basic constituents of the nuclei, lamins are assembled in a filamentous meshwork, providing structural stability. Importantly, only after nanobody production in an *E. coli* strain, engineered to promote disulfide bond formation in the cytosol, and application of an optimized coupling procedure, specific labeling of the endogenous nuclear lamina was facilitated with α-Lamin Fbs equipped with sCy5 or ATTO655 at labeling ratios of 0.8 to 1.0, respectively. Noteworthy, different procedures led to successful dye conjugation, but were accompanied with loss in specificity (**Figure 1 – Figure Supplement 4**). Similar results were obtained if lysine residues on the nanobody surface were stochastically modified via *N*-hydroxysuccinimide (NHS) Alexa647 (**Figure 1 – Figure Supplement 4a**). Contrary, the α-GFP Fb always retained high target specificity in fixed cells, independent of all described routes for production and labeling (**Figure 1 – Figure Supplement 5**). Since the highest signal-to-background ratio was achieved after dye-conjugation to the N-terminal cysteine of the α-Lamin Fb (**Figure 1f,g**; **Supplementary Video 1** and **Figure 1 – Figure Supplement Fig. 4**), labeling at this position was employed for all further experiments.

### Live-cell protein labeling at nanomolar fluorobody concentrations

Once the specificity in fixed cells was proven, we aimed at live-cell protein labeling. For the fast and efficient Fb transfer, we made use of high-throughput, microfluidic cell squeezing. Here, transient pore formation of the plasma membrane is induced by forcing cells through micrometer constrictions of microfluidic devices (**Figure 2 – Supplement Figure 1**). This approach has shown to bypass endosomal uptake and offered precise control of the transduced cargo concentration combined with high-throughput delivery (1,000,000 cells/s) and minimal cytotoxicity (< 10%).^23,24^ Initially, α-GFP Fbs were delivered into HeLa Kyoto cells expressing the three selected FP-tagged proteins and the intracellular Fb distribution was analyzed 1 to 3 h after cell transfer. In all cases, the fluorescence of the α-GFP Fb showed a high degree of co-localization with FP-tagged H2B, TAP1, or lamin A, reflected by Pearson’s coefficients ranging from 0.864 to 0.911 (**Figure 2a** and **Figure 2 – Supplement Figure 2a**). Exemplary, an excellent correlation between the fluorescence intensity profiles of TAP1-mVenus and the α-GFP^sCy3^ Fb was observed (**Figure 2b,c**; Pearson’s coefficient of 0.926). Notably, all transduced cells displayed a very low background staining and the intensity of the Fb reflected the variability in the POI expression level. We observed an increased Fb binding in cells with high levels of FP-tagged POIs, whereas cells expressing low amounts of the POIs displayed weaker fluorescence intensities (**Figure 2 – Supplement Figure 3**). One explanation for this correlation is found in the release of unbound binders during membrane resealing as well as in the nanomolar concentration of Fbs in the squeezing buffer, both synergistically contributing to a high signal-to-background staining (**Figure 2d**).^22^ Indeed, untransfected cells indicated a very low level of uniformly intracellular distributed Fbs (**Figure 2 – Supplement Figure 4**). This is in line with the localization of the α-GFP-mCherry chromobody or mCherry expressed in HeLa Kyoto cells (**Figure 2 – Supplement Figure 5a,b**). In contrast, expression of α-GFP-mCherry chromobodies together with FP-tagged POIs often resulted in overabundance of the binder due to mismatched expression levels (**Figure 2 – Supplement Figure 5c-e**). Here, a high signal-to-background labeling is impaired in cells with high expression levels of chromobodies, while very low levels entail high sensitivity to photobleaching. These drawbacks were successfully abolished by microfluidic Fb transfer. To test the long-term stability and persistence of Fb labeling, cells were imaged 20 h after transduction. Remarkably, the POIs were still decorated by α-GFP Fbs, highlighting their long-term specificity and in-cell robustness (**Figure 2 – Supplement Figure 2**). Over time, however, cytosolic punctae and a higher background signal were detected, indicating incipient Fb degradation. Collectively, the adaptive, fine-tuned intracellular delivery of Fbs by cell squeezing promoted low-background labeling of genetically tagged proteins.

**Figure 2.**
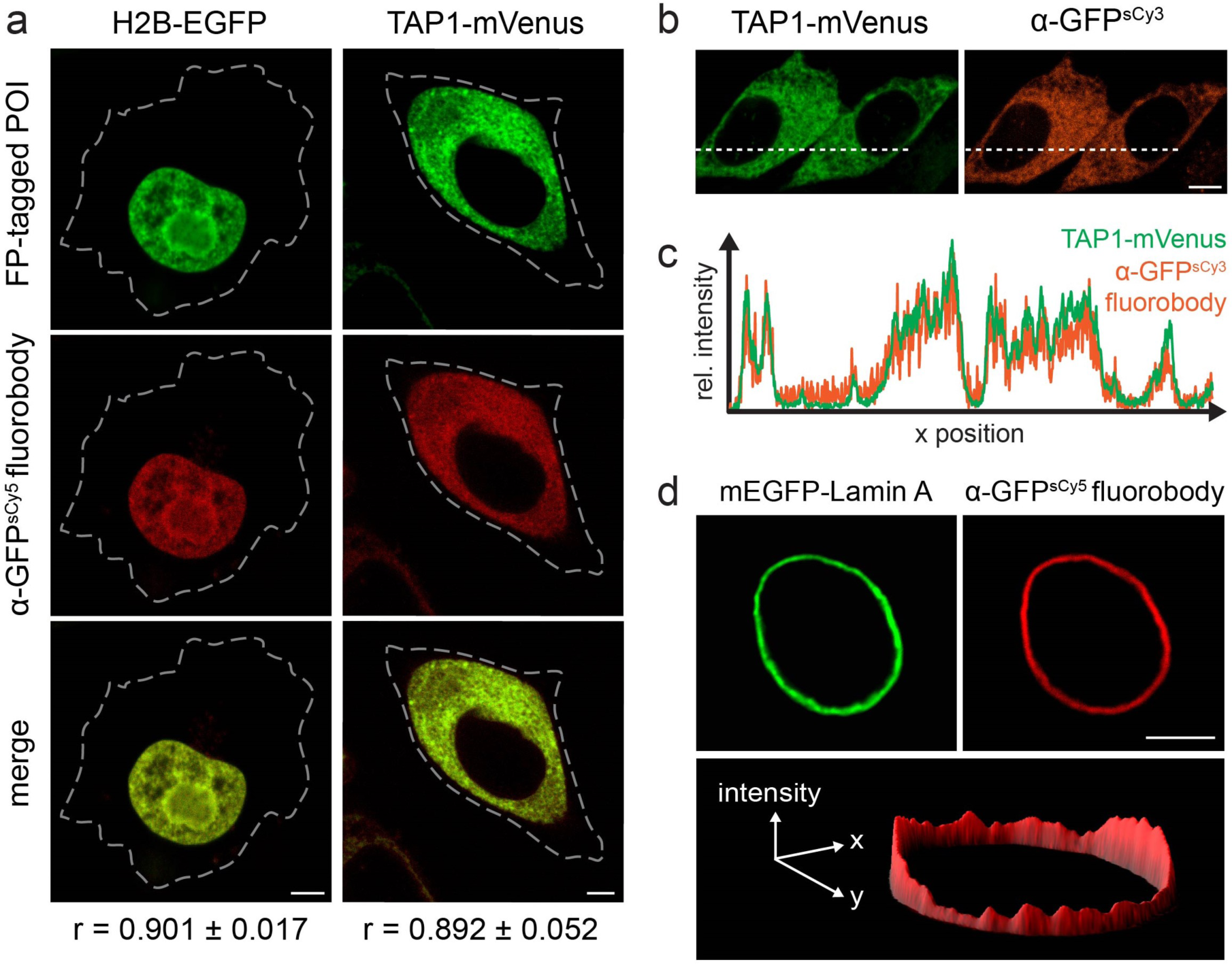
Ι Live-cell protein labeling with high-performance fluorobodies. (**a**) Protein labeling in living cells at different subcellular localizations. The α-GFP^sCy5^ Fb (200 nM, red) was delivered via cell squeezing into HeLa Kyoto cells, expressing H2B-EGFP or TAP1-mVenus (green). Excellent co-localization (merge channel) of the Fbs with both FP-tagged POIs was observed 3 h post squeezing by CLSM. The Pearson’s coefficients (r, bottom) ranging from 0.864 to 0.905 were calculated from 8-10 individual cells. Dashed lines indicate cell borders. (**b**) Live-cell visualization of TAP1-mVenus (green) by α-GFP^sCy3^ Fb (orange) 20 h after squeezing. Cross-sections of relative fluorescence intensity profiles of TAP1-mVenus and α-GFP^sCy3^ (along dashed line) showed specific and low-background labeling in live cells by the α-GFP Fb (**c**). Pearson’s coefficient: 0.926. (**d**) 3D intensity plot of a representative HeLa Kyoto cell expressing mEGFP-Lamin A (green) was stained by α-GFP^sCy5^ Fb (red) and showed a very high signal-to background ratio. All images were taken by CLSM. Scale bar: 5 µm.

### High signal-to-background tracing of endogenous proteins in live mammalian cells

We next focused on live-cell labeling of endogenous protein networks. With regard to their extenuated abundance, the utilization of high-performance fluorescent or fluorogenic probes is essential to enhance sensitivity. We selected two intermediate filaments as targets. Specifically, we used the engineered α-Lamin Fb to visualize the native lamin meshwork of the nuclear envelope and α-Vimentin^ATTO488^ for tracing vimentin as a cytoskeletal component, crucial for organelle positioning. Target specificity of both Fbs was verified beforehand in fixed cells (**Figure 1f,g** and **Figure 3 – Supplement Figure 1**). In live HeLa Kyoto cells expressing α-Lamin-EGFP chromobody, decoration of the nuclear lamina was accompanied with high background staining and cytosolic punctae, presumably due to saturation of the targeting machinery (**Figure 3 – Supplement Figure 2**). Contrarily, specific decoration of endogenous lamin with a concomitant negligible background was visualized 3 h after cell squeezing utilizing the α-Lamin^ATTO655^ Fb (**Figure 3a**). To the best of our knowledge this constitutes the first labeling of an endogenous protein inside living cells by Fbs. Notably, even 20 h after cell squeezing, labeling of the nuclear lamina was still persistent. Nevertheless, higher background, mainly as cytosolic punctae, was observed (**Figure 3 – Supplement Figure 3**), in line with results obtained for the α-GFP Fb (**Figure 2 – Supplement Figure 2b**). Of note, specific targeting was unaffected by the utilized fluorophore (ATTO655 or sCy5; **Figure 3a** and **Figure 3 – Supplement Figure 3**). To further broaden the range of applications, α-Vimentin^ATTO488^ was delivered into cells. The well-defined filamentous structure of endogenous vimentin was visualized with a high signal-to-background ratio 3 h as well 20 h after cell squeezing (**Figure 3b,c** and **Figure 3 – Supplement Figure 4**).

**Figure 3.**
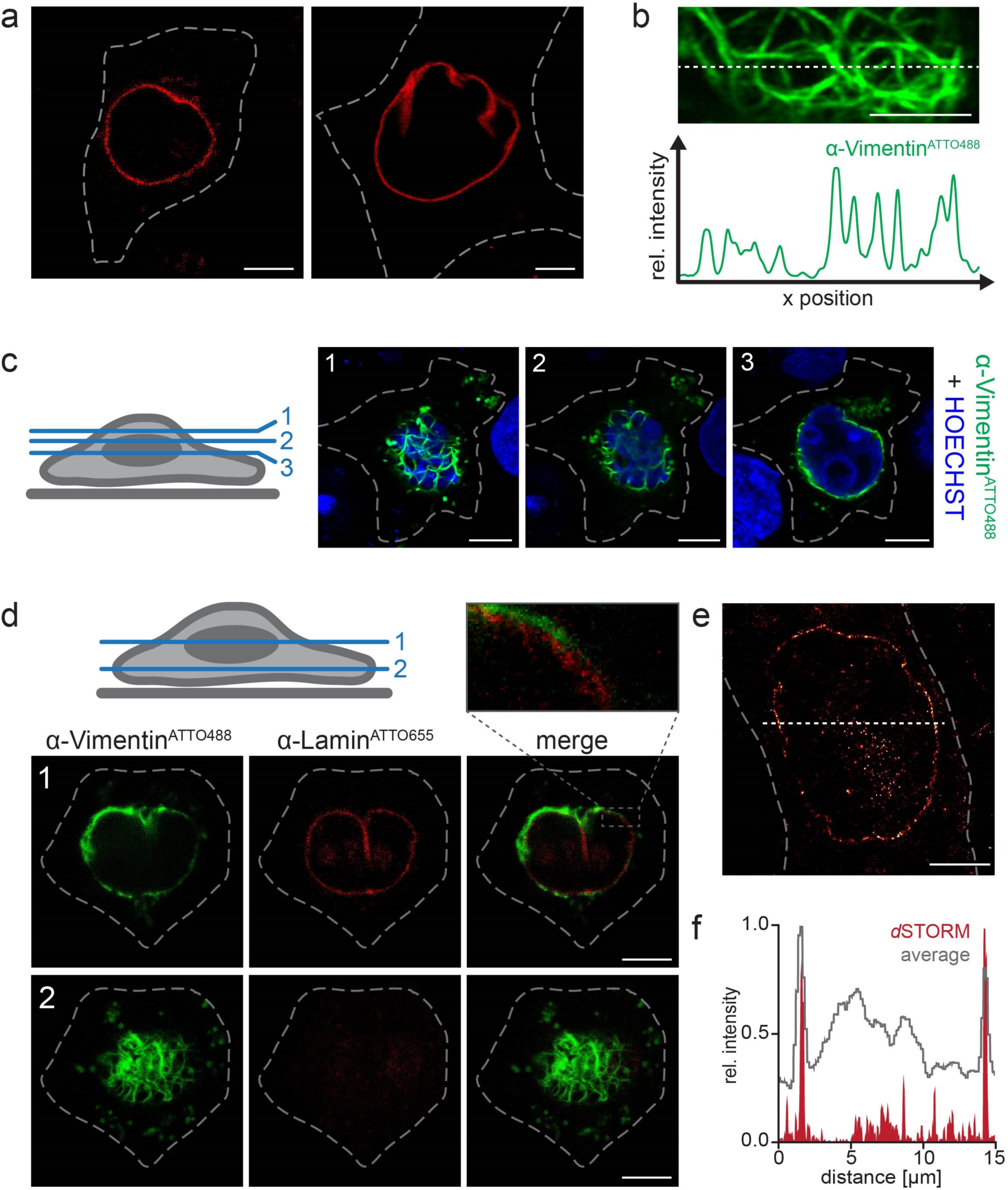
Ι Dual-color live-cell tracing of endogenous targets and live-cell super-resolution imaging by fluorobodies. (**a**) The α-Lamin^ATTO655^ Fb (200 nM) was transferred into HeLa Kyoto cells via squeezing. Confocal imaging after 3 h showed specific labeling of the native nuclear lamina in living cells with high signal-to-background ratio. (**b**) HeLa Kyoto cells were squeezed with 50 µg/ml of α-Vimentin^ATTO488^. The Fb decorated vimentin network was explicitly traced 3 h post squeezing and visualized as a dense meshwork around the nucleus (HOECHST-positive, blue) in different confocal z-planes (1-3; indicated in scheme, left). (**c**) Fluorescence intensity cross-section of a live HeLa Kyoto cell transduced with α-Vimentin^ATTO488^ (along dashed line) showed specific and distinct decoration of the endogenous vimentin meshwork with high signal-to-background ratio. The respective network was localized beneath the nucleus (corresponding to z-plane 2 in d). (**d**) Dual-color labeling of endogenous lamin and vimentin in living cells. α-Lamin^ATTO655^ and α-Vimentin^ATTO488^ were simultaneously transferred into HeLa Kyoto cells. 3 h after cell squeezing, spatially distinct labeling of endogenous lamin (red) and vimentin (green) was visualized by multiplexed imaging. Magnification (box, top) of the α-Vimentin^ATTO488^ stained vimentin network showed the three-dimensional organization around the nucleus. The respective confocal z-planes are indicated in the scheme. (**e**) Reconstructed *d*STORM image of endogenous lamin specifically labeled with α-Lamin^ATTO655^ Fb (5 h after squeezing). The sensitive and high degree of Fb labeling allowed for visualization of the native nuclear envelope with nanometer precision. (**f**) Comparison of *d*STORM analysis with the averaged fluorescence intensity. Cross section of the intensity profiles (along dashed line in e) displayed the enhanced resolution by *d*STROM. Images were taken by CLSM (a-d) with the Airy detector (except for d) or *d*STORM (e). Dashed lines indicate the cell border. Scale bars: 5 µm.

### Multiplex and super-resolution analysis of native cellular protein networks

Encouraged by these results, we performed multi-color live-cell labeling by combining both binders. 3D mapping of cell topology via multiplexed imaging revealed simultaneous and specific decoration of the nuclear lamina (**Figure 3d**, red) and vimentin filaments (**Figure 3d**, green) at endogenous level, hardly realized by multiple knock-ins utilizing the CRISPR/Cas9 technology. The spatial distinct distribution allowed discriminating between both structures by live-cell immunofluorescence imaging. In combination with their precisely adjusted delivery by cell squeezing, the small Fbs, site-specifically labeled with minimal and bright fluorescent dyes (≤ 1 nm) provides an excellent showcase for live-cell super-resolution microscopy. Consequently, α-Lamin^ATTO655^ Fb transduced cells were subjected to *direct* stochastic optical reconstruction microscopy (*d*STORM).^28^ We obtained images of the endogenous nuclear lamina in living cells with an average resolution of ~ 55 nm (localization precision of 23.4 ± 8.3 nm; **Figure 3e**) and significantly increased contrast compared to wide-field imaging (**Figure 3f**). Strikingly, *d*STORM analysis of α-Lamin^ATTO655^ Fb in fixed cells yielded virtually identical average localization precision, but an increased signal-to-background-ratio was obtained in living cells, devoid of fixation artifacts (**Figure 3 – Supplement Figure 5**).

## Discussion

In general, the much lower level of endogenous proteins makes their specific and low-background tracing very challenging. To circumvent this constrain, we established a method for live-cell visualization of endogenous proteins at nanometer resolution and with minimal disturbance and high reproducibility (n ≥ 3 for every POI). This robust and generic technique is versatile in the choice of the nanobody and bright organic fluorophore and allows for low-background tracing of high as well as low abundant native protein networks in living cells. In case of our selected target structures, Fb decoration was persistent for several hours after Fb transfer. However, increased background after 20 h suggests that long-term protein tracing can be affected. This obstacle is experimentally manageable, depending on the turnover rate of the POI as well as the rate of cell division. Alternative approaches to tag endogenous proteins such as the generation of stable cell lines utilizing *e.g.* CRISPR/Cas9 for low chromobody levels are highly promising, yet very labor-intensive and hardly realizable for multiple knock-ins.

Collectively, our devised strategy strikes a new path for the live-cell visualization of endogenous targets by high quantum yield fluorobodies to study their precise intracellular localization and trafficking. The simplicity and robustness of the methodology can be extended to deliver multiple probes simultaneously for multiplexed imaging and has tremendous potential for other types of microscopy (*e.g.* electron microscopy). Even the Fb targeting of endogenous proteins in embryonic stem cells or patient-derived primary immune cells is technically feasible via cell squeezing. Moreover, our method provides access of exogenously derived high-affinity binding scaffolds (*e.g.* fibronectin-based binding proteins or designed ankyrin repeat proteins) for 3D and multi-color super-resolution imaging in native cells and opens avenues to label cellular ultrastructures at endogenous level for correlated light and electron microscopy (CLEM) or lattice light-sheet microscopy. Overall, the readily and feasible expansion of the methodology will pioneer the non-genetically encoded visualization of cellular networks and subcellular structures and hence provide complementary and unique information at unprecedented precision. Thus, fluorobody delivery by cell squeezing for in-cell tracing will likely find applications supporting routine imaging of multiple native cellular assemblies with advanced imaging techniques.

## Acknowledgement

The German Research Foundation (Cluster of Excellence EXC 115 – Macromolecular Complexes to R.W., M.H, and R.T.; SPP 1623, RTG 1986 to R.T. and CRC 807 to M.H. and R.T.) and the Volkswagen Foundation (A117158 to M.H., R.W., and R.T.) supported this work. We thank Dr. Eric Geertsma (Goethe University Frankfurt, Germany) for providing the sequence coding for the α-GFP nanobody and helpful discussions. We thank Dr. Heinrich Leonhardt (LMU Munich, Germany) for generously providing the plasmid coding for α-Lamin-EGFP.

## Authors Contribution

A.K. performed the nanobody labeling and cell squeezing experiments. S.H., A.K., and F.T. carried out the protein expression and purification. A.R. and M.H. performed the *d*STORM imaging and analysis. A.K., R.W., and R.T. wrote the manuscript and analyzed the data. R.W. and R.T. conceived the ideas and directed the work.

## Methods

### Plasmid construction

For expression in *E. coli*, the α-GFP nanobody,^26^ equipped with a C-terminal His_6_-tag for purification via metal ion affinity chromatography, was inserted into the pET21a(+) plasmid. For site-specific labeling, a free single cysteine was introduced C-terminally to the His_6_-tag. The α-GFP nanobody was PCR amplified by the following primers: fw 5’-GCGCGC<u>AAGCTT</u>*AAGGAG*ATATACAT**ATG**CAGGTTCAGCTGGTTGAAAGCGGTGGTGCAC-3’ (Hind III restriction site underlined, RBS italic, start codon bold); rev 5’-GCGC<u>CTCGAG</u>G**TCA<u>GCA</u>***GTGGTGATGGTGATGATG*GCTGCTAACGGTAACCTGGGTGCCC-3’ (XhoI restriction site underlined, stop codon bold, cysteine bold and underlined, His_6_-tag italic). For expression in mammalian cells, α-GFP as well as mCherry were PCR amplified and introduced into the pcDNA3.1(+) plasmid, resulting in a plasmid coding for α-GFP^mCherry^. fw 5’-GCGCGC<u>GCTAGC</u>ACC**ATG**CAGGTTCAGCTGGTTGAAAGCGG-3’ (NheI restriction site underlined, start codon bold) and rev 5’-GCGCGC<u>GGTACC</u>GCTGCTAACGGTAACCTGGGTGCCC-3’ (Acc65I restriction site underlined) for the α-GFP nanobody and fw 5’-CGCGC<u>GGTACC</u>**ATG**GTGAGCAAGGGCGAGGAGCTGTTC-3’ (Acc65I restriction site underlined, start codon bold) and rev 5’-GCGCGC<u>CTCGAG</u>**TTA**CTTGTACAGCTCGTCCATGCCGAGAG-3’ (XhoI restriction site underlined, stop codon bold) for mCherry. The α-Lamin nanobody was also inserted into the pET21a(+) plasmid and equipped with a His_6_-tag for purification. Using α-Lamin-EGFP as PCR template, α-Lamin was amplified, introducing a C-terminal cysteine (α-Lamin^His6-Cys^), a N-terminal cysteine (^Cys^α-Lamin^His6^), or replacing serine 9 by a cysteine (α-Lamin^S9C-His6^). α-Lamin^His6-Cys^: fw 5’-GCGC<u>AAGCTT</u>*AAGGAGA*TATACAT**ATG**GCTCAGGTACAGCTGCAGGAGTCTGGAGGAGG-3’ (HindIII restriction site underlined, start codon bold, RBS italic); rev 5’-GCGC<u>CTCGAG</u>**TTAACA***GTGGTGATGGTGATGATG*CGATATCGAATTCCTTGAGGAGACGGTGACCTGGG-3’ (XhoI restriction site underlined, stop codon bold, cysteine bold and underlined, His_6_-tag italic). To attach the PelB leader sequence and the cysteine, ^Cys^α-Lamin^His6^ were amplified in consecutive extending PCRs, each of them being the template for the subsequent amplification. The rev primer 5’-GCGCGC<u>CTCGAG</u>**TTA***GTGGTGATGGTGATGATG*GGTGGCGACCGGCCG-3’ (XhoI restriction site underlined, stop codon bold, His_6_-tag italic) was thereby applied in combination with the following fw primers. ^Cys^α-Lamin^His6^: fw1 5’-*GGATTGTTATTACTCGCGGC CCAGCCGGCC***TGT**GCTCAGGTACAGCTGCAGGAGTCTGGAGGAGG-3’ (cysteine bold, part of pelB leader sequence italic), fw2 5’-GCGC<u>GCTAGC</u>**ATG***AAATACCTATTGCCTACGGCAGCCGCTGGATTGTTATTACTCGCGGCCCAGCCGGCC*-3’ (NheI restriction site underlined, start codon bold, pelB leader sequence italic). α-Lamin^S9C-His6^: fw1 5’-ATGGCTCAGGTACAGCTGCAGGAGTGTGGAGGAGGCTTGGTGCAGGCAGGGGGGTCTCTG-3’, fw2 5’-*GGATTGTTAT TACTCGCGGCCCAGCCGGCC*ATG GCTCAGGTACAGCTGCAGGAG**TGT**GGA-3’ (cysteine bold, part of pelB leader sequence italic), fw3 5’-GCGC<u>GCTAGC</u>**ATG***AAATACCTATTGCCTACGGCAGCCGCTGGATTGTTATTACTCGCGGCCCAGCCGGCC*-3’ (NheI restriction site underlined, start codon bold, pelB leader sequence italic). The plasmid coding for H2B-EGFP was obtained from Addgene (plasmid 11680). The generation of plasmids coding for core TAP1 (TAP1-mVenus) and mEGFP-Lamin A was described previously.^23,29^

### Nanobody production

The nanobodies were produced in *E. coli* BL21 (DE3, Life technologies) or *E. coli* SHuffle T7 Express (New England Biolabs), whereas the latter contains a chromosomal copy of a disulfide isomerase, engineered to produce proteins with disulfide bonds in the cytosol. Both *E. coli* strains were transformed with the vectors, coding for α-GFP and α-Lamin nanobodies with engineered cysteines. A single colony was inoculated in LB medium containing 100 µg/ml ampicillin and 2% (w/v) glucose. After overnight incubation (37 °C for *E. coli* BL21 (DE3) and 30 °C for *E. coli* SHuffle T7 Express, 180 rpm), 1 l Lysogeny Broth (LB) medium with 100 µg/ml ampicillin was inoculated and protein expression was induced at A_600_ of 0.7-0.8 by 1 mM isopropyl-β-thiogalactopyranoside (IPTG, Sigma-Aldrich) and conducted for 5-20 h (30 °C, 180 rpm). After centrifugation (20 min, 4°C, 5,000x g), the cell pellets were directly used for protein purification or stored at -20 °C.

### Protein purification

Two different protocols were used for purification. For α-GFP^His6-Cys^ and α-Lamin^His6-Cys^ expressed in *E. coli* BL21 (DE3), cell pellets were resuspended in lysis buffer (20 mM Tris pH 8.0, 100 mM NaCl, 1 mM phenylmethylsulfonyl fluoride (PMSF), 0.5 mM β-mercaptoethanol (β-ME)), followed by sonication for cell lysis. Cell debris were pelleted by ultracentrifugation (100,000x g, 30 min, 4 °C). For metal affinity chromatography, the supernatant was incubated with 3 ml Ni-NTA Sepharose beads (GE Healthcare) in the presence of imidazole (15 mM). Afterwards, the beads were washed in purification buffer (50 mM Tris pH 8.0, 300 mM NaCl, 50 mM imidazole, 0.5 mM β-ME). His-tagged proteins were eluted by 200 mM imidazole/purification buffer. Protein concentration was determined at A_280_. Buffer exchange to phosphate-buffered saline (PBS) containing 0.5 mM β-ME by Zeba spin desalting columns (7K) was followed by concentration via Amicon Ultra Filters (3 K). β-ME was directly removed before labeling the nanobodies using Zeba spin desalting columns (7K). Alternatively, nanobodies expressed in *E. coli* T7 Express were purified as described above, but at pH 7.5 and without β-ME.

### Protein modification

Fluorescent dyes were either attached to the engineered cysteines or to lysines (only for α-Lamin^His6-Cys^). The engineered cysteines of nanobodies stored in PBS without β-ME were reduced by incubation with 15 mM TCEP for 10 min on ice and subsequent buffer exchange by Zeba spin desalting columns (7K). Cysteine labeling was conducted following two different protocols. On the one hand, nanobodies were incubated with sulfo-Cy5 (sCy5) or sulfo-Cy3 (sCy3) maleimide in PBS (pH 7.4) for 1.5 h at 4 °C. Labeling was conducted at a protein concentration of approximately 1 mg/ml with 1.2-fold molar excess of the dye. For the alternative labeling protocol, buffer was exchanged to maleimide labeling buffer after TCEP reduction (100 mM KH_2_PO_4_ pH 6.4, 150 mM NaCl, 1 mM EDTA, 250 mM sucrose). Afterwards, the fluorophore was added in a 1.2- fold molar excess, immediately followed by neutralization to pH 7.5 using K_2_HPO_4_ and 1.5 h incubation at 4 °C. For lysine labeling, nanobodies in PBS (pH 7.4) were mixed 1:20 with 0.1 mM NaHCO_3_ (pH 9.0) to achieve pH 8.3 for labeling. Next, Alexa647 NHS ester was added with a 5-fold molar excess. Labeling was performed for 1 h at 20 °C before buffer exchange and removal of excess dye with Zeba spin desalting columns (7K, Thermo Scientific). Fluorescently labeled nanobodies were analyzed by SDS-PAGE and in-gel fluorescence (Typhoon 9400 Imager; l_ex/em_ 630/670 nm). Fbs were finally purified by size-exclusion chromatography in PBS using a KW404-4F column (Shodex; flow rate 0.3 ml/min). All labeled nanobodies were stored in PBS at 4 °C. The nanobody-to-fluorophore ratio was determined using absorption (A_280_ for nanobodies, A_550_ for sCy3, A_663_ for ATTO655, A_649_ for sCy5) and their respective extinction coefficient ε (0.22·10^5^ M^−1^·cm^−1^ for α-Lamin Nb, 0.27·10^5^ M^−1^·cm^−1^ for α-GFP Nb, 1.36·10^5^ M^−1^·cm^−1^ for sCy3, 1.25·10^5^ M^−1^·cm^−1^ for ATTO655, and 2.50·10^5^ M^−1^·cm+for sCy5).

### Cell culture and transfection

HeLa Kyoto cell lines were cultivated in Dulbecco’s Modified Eagle Medium (DMEM) with 4.5 g/l glucose (Gibco) containing 10% (v/v) fetal calf serum (FCS, Gibco). Cells were regularly tested for Mycoplasma contamination.^30^ Cells were transiently transfected using Lipofectamine 2000 (Life technologies), following the manufacturer’s instructions. For staining of fixed cells, 2.10^4^ cells per well were seeded into 8-well on cover glass II slides (Sarstedt) and transfected at ~ 80% confluency. For cell squeezing, 8.10^5^ cells were seeded into 6-well cell culture plates (Greiner) and transiently transfected. After transfection, cells were incubated for 12-24 h at standard cell culture conditions until experiments were performed.

### Cell fixation and labeling

Cells were washed with PBS (Sigma-Aldrich), fixed with 4% formaldehyde (Roth)/PBS for 10-20 min at 20 °C and permeabilized using 0.1% Triton X-100 (Roth)/PBS (10 min, 20 °C). After blocking with 5% (w/v) bovine serum albumin (BSA; Albumin Fraction V, Roth) in PBS for 1 h at 20 °C, cells were stained. Staining was performed with fluorobodies (diluted in 1% BSA/PBS) at concentrations ranging from 50 to 500 nM (indicated in each case). Subsequently, cells were washed with PBS and optionally stained with 0.1 µg/ml 4',6-diamidino-2-phenylindole (DAPI, Sigma-Aldrich) in 1% BSA/PBS for 30 min to 1 h at 20 °C. After washing with 5% BSA/PBS (3x), cells were post-fixed with 2% formaldehyde/PBS (15 min, 20 °C) and stored in PBS until imaging was performed.

### Confocal laser scanning microscopy

Imaging was performed using the confocal laser scanning (LSM) microscope LSM880 (Zeiss) or TCS SP5 microscope (Leica) with Plan-Apochromat 63x/1.4 Oil DIC objectives. To avoid crosstalk between different fluorophores, images were acquired sequentially. The following laser lines were used for excitation: 405 nm for DAPI; 488 nm for EGFP, mEGFP, and mVenus; 565 nm for sCy3; 633 nm for sCy5 and ATTO655. Live-cell imaging of endogenous proteins (except for dual-color imaging with ATTO488 and ATTO655) was conducted with the Airy scan detector of the LSM880 in the SR-mode. For 3D reconstructions of labeled protein networks, z-stack were recorded with the Airy scan detector (distance between stacks ~ 200 nm). For live-cell imaging, an incubation system was used to keep cells at 37 °C and supply them with 5% CO_2_ during imaging. ImageJ^31^, Fiji^32^, and Zen 2.3 black (Zeiss) were used for image analysis and to analyze Pearson’s coefficients from 8-11 individual cells per condition. The mean and standard deviation were calculated.

### Fluorobody transfer by cell squeezing

Squeezing was performed using a chip with constrictions of 7 µm in diameter and 10 µm in length (CellSqueeze 10-(7)x1, SQZbiotech). In all microfluidic experiments, a cell density of 1.5.10^6^ cells/ml in 10% (v/v) FCS/PBS was squeezed through the chip at a pressure of 30 psi. Transduction was conducted at 4 °C to block cargo uptake by endocytosis.^23^ During squeezing, the following Fb concentrations in the surrounding buffer were used: 100-200 nM of α-GFP, 200-500 nM of α-Lamin, and 50 µg/ml of α-Vimentin. After squeezing, cells were incubated for 5 min at 4 °C to reseal the plasma membrane. Squeezed cells were washed with DMEM containing 10% FCS, seeded into 8-well on cover glass II slides (Sarstedt) or collagen-coated dishes (µ-Slide 8 Well, Collagen IV coverslip, ibidi) in DMEM containing 10% FCS, and cultured at 37 °C and 5% CO_2_. Confocal imaging was performed 1 h and 20 h after squeezing. Squeezing experiments for live-cell protein labeling highly reproducible in at least three independent experiments for every POI.

### Super-resolution microscopy

For super-resolution imaging of the α-Lamin^ATTO655^ Fb in live cells, a custom-built microscope was used. Samples were illuminated with 643 nm (iBeam smart, Toptica Photonics) laser beams and a 405 nm reactivation laser (Cube 405-50C, Coherent) in total internal reflection fluorescence (TIRF) mode. The excitation light was focused on the back focal plane of a 100x oil objective (Plan Apo 100x TIRFM, NA 1.45, Olympus), mounted on an inverted microscope (Olympus IX71). The emission was recorded using an EMCCD camera (IxonUltra, Andor) with frame-transfer mode, 1x pre-amplifier gain and EM gain set to 200. For every sample, 40,000 images were recorded at a frame rate of 33 Hz. Image reconstruction was performed with rapidSTORM^33^ using the smooth by differences of averages option with a signal-to-noise ratio of 20, fitting only PSFs with FWHM ranging from 240-520 nm and X-Y-ratio of 0.7-1.3. The average localization precision (σ_loc_) defined as the nearest neighbor distance in adjacent frames^34^ was calculated to 23.4 ± 8.3 nm for living and 23.2 ± 4.6 nm for fixed cells (mean and standard deviation of three individually analyzed cells, respectively) using LAMA software.^35^ The average image used for visualization of nuclear intensity profiles in Figure 3f was generated by averaging the fluorescence intensity over 100 frames and subsequent background subtraction with a rolling ball method (20 pixel radius) in ImageJ and Fiji.

## Data availability

The datasets generated during and/or analyzed during the current study are available from the corresponding author on reasonable request.

## SUPPLEMENTAL FIGURES

**Figure 1 – Figure Supplement 1.**
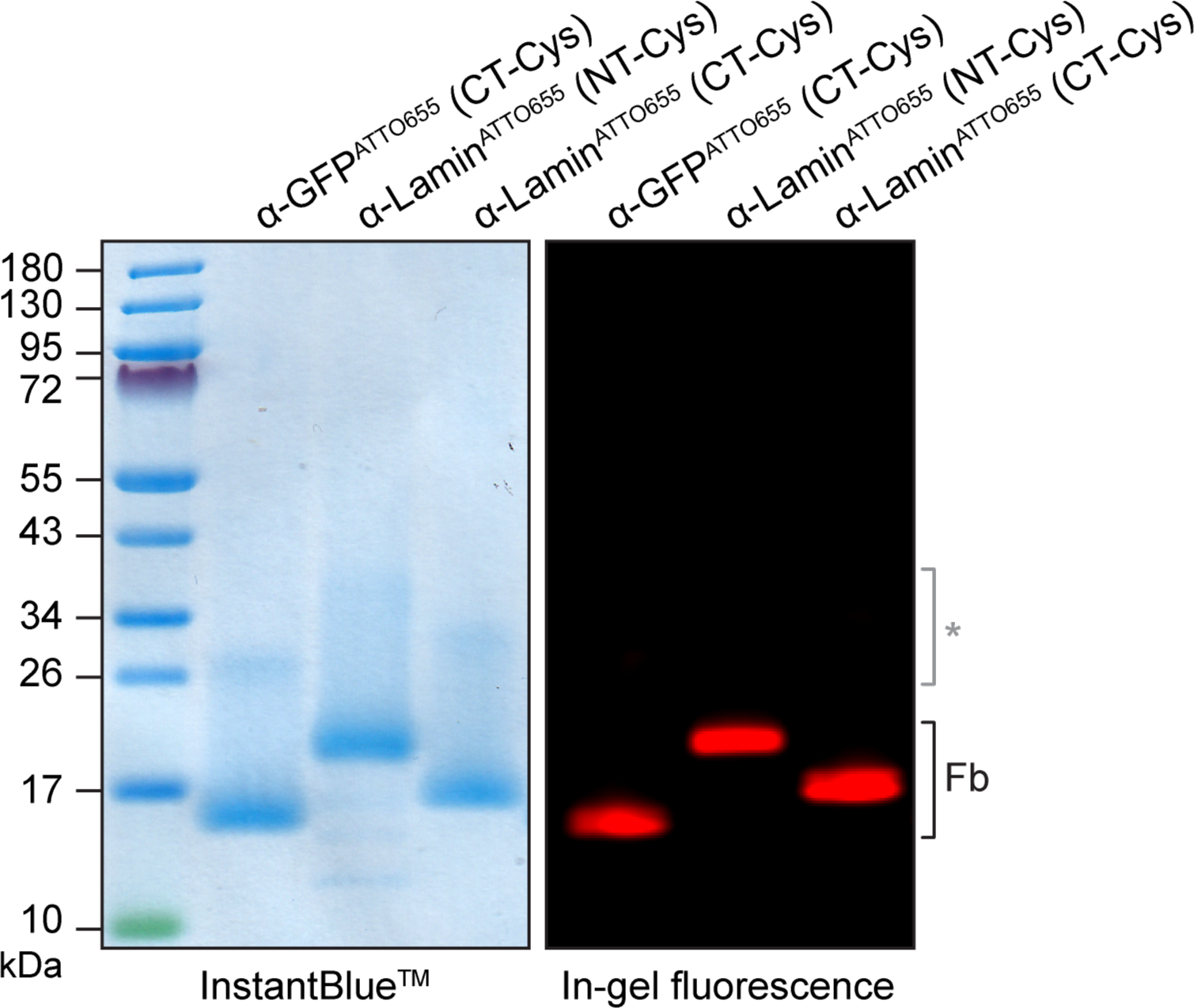
Site-specific fluorescent labeling of nanobodies. Single engineered surface cysteines of nanobodies were modified with maleimide ATTO655. SDS-PAGE analysis showed covalent labeling of the engineered Fbs. Detection was performed by InstantBlue^TM^ staining and by in-gel fluorescence with λ_ex/em_ 630/670 nm. A minor portion of Fb dimers (*) was observed. The higher molecular weight of α-Lamin (NT-Cys) is due to an elongated linker between Nb and C-terminal His_6_-tag.

**Figure 1 – Figure Supplement 2.**
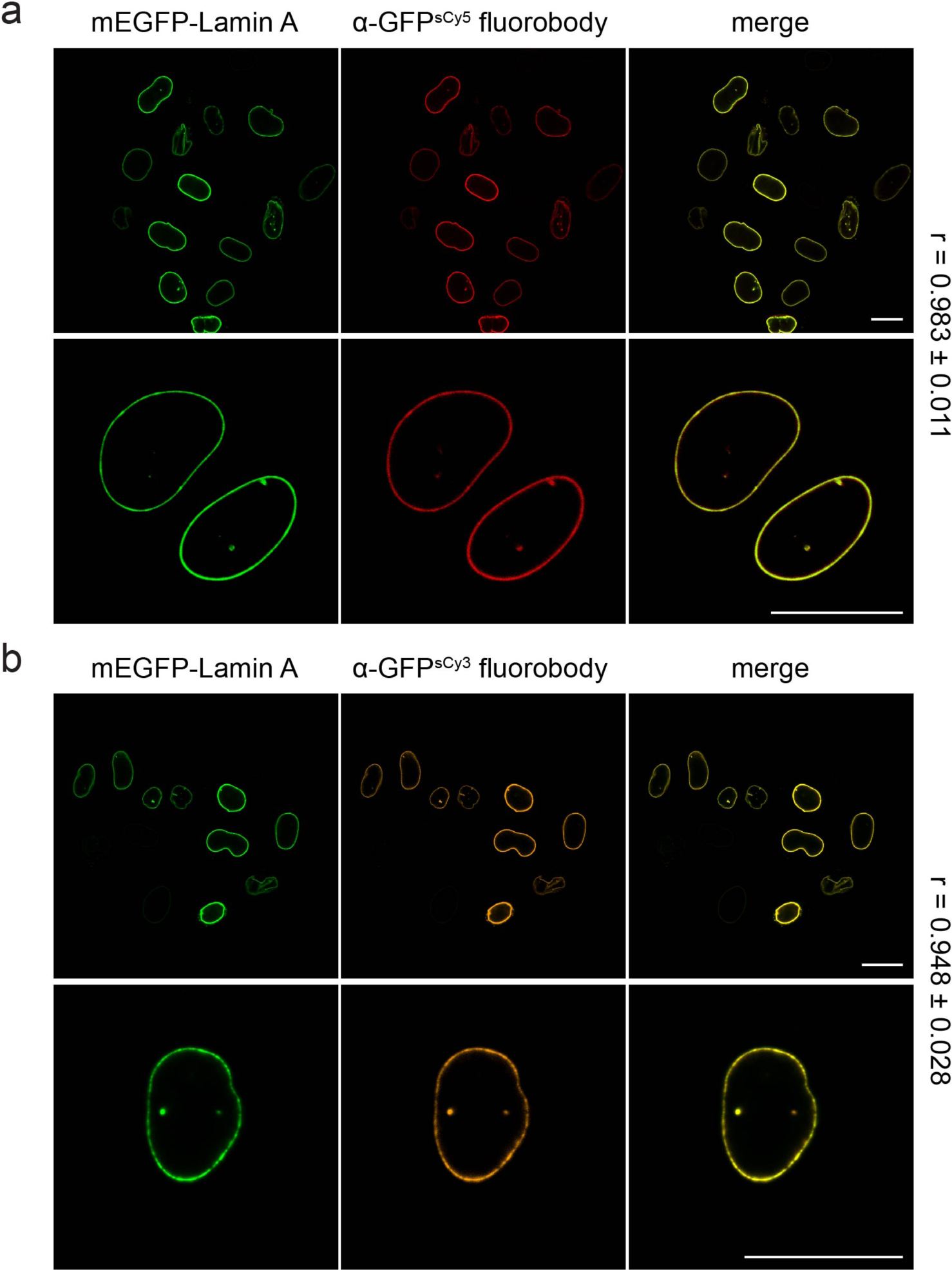
Specific labeling of mEGFP-Lamin A with α-GFP^sCy3^ and α-GFP^sCy5^ fluorobodies. After fixation and permeabilization, HeLa Kyoto cells expressing mEGFP-Lamin A were labeled with 100 nM of α-GFP^sCy5^ Fb (**a**) or α-GFP^sCy3^ Fb (**b**) and analyzed *via* CLSM. mEGFP-Lamin A was specifically labeled by the Fbs with very low background and the staining intensity correlated with the POI´s expression level. Scale bars: 20 µm.

**Figure 1 – Figure Supplement 3.**
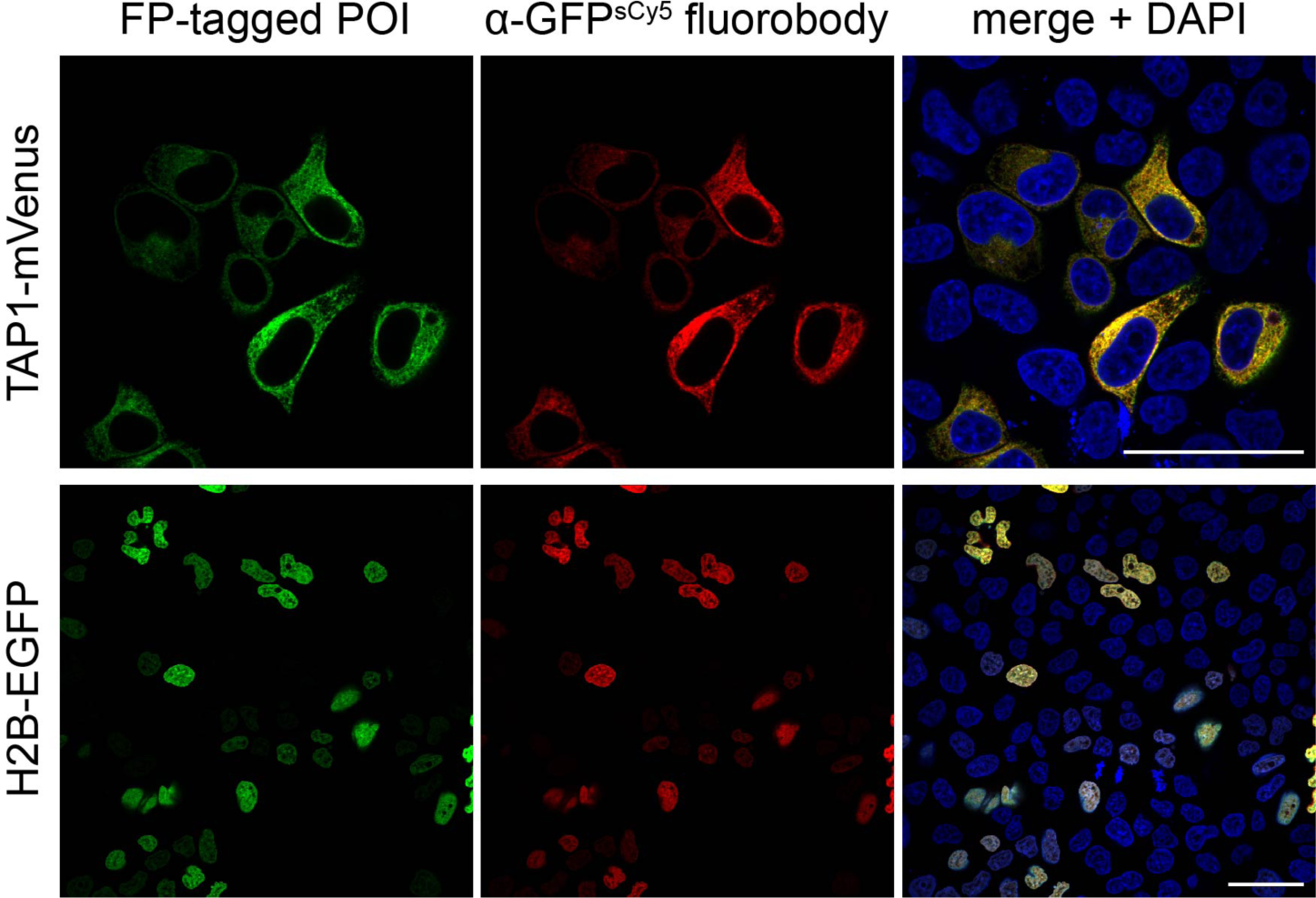
Correlation of the target protein expression level with the labeling density of the α-GFP fluorobody. After fixation and permeabilization, HeLa Kyoto cells expressing TAP1-mVenus or H2B-EGFP were stained with α-GFP^sCy5^ Fb (50 nM). The labeling intensity highly correlates with the expression level of both FP-fused POIs, reflecting stoichiometric target decoration. Imaging was performed by CLSM. Scale bars: 50 µm.

**Figure 1 – Figure Supplement 4.**
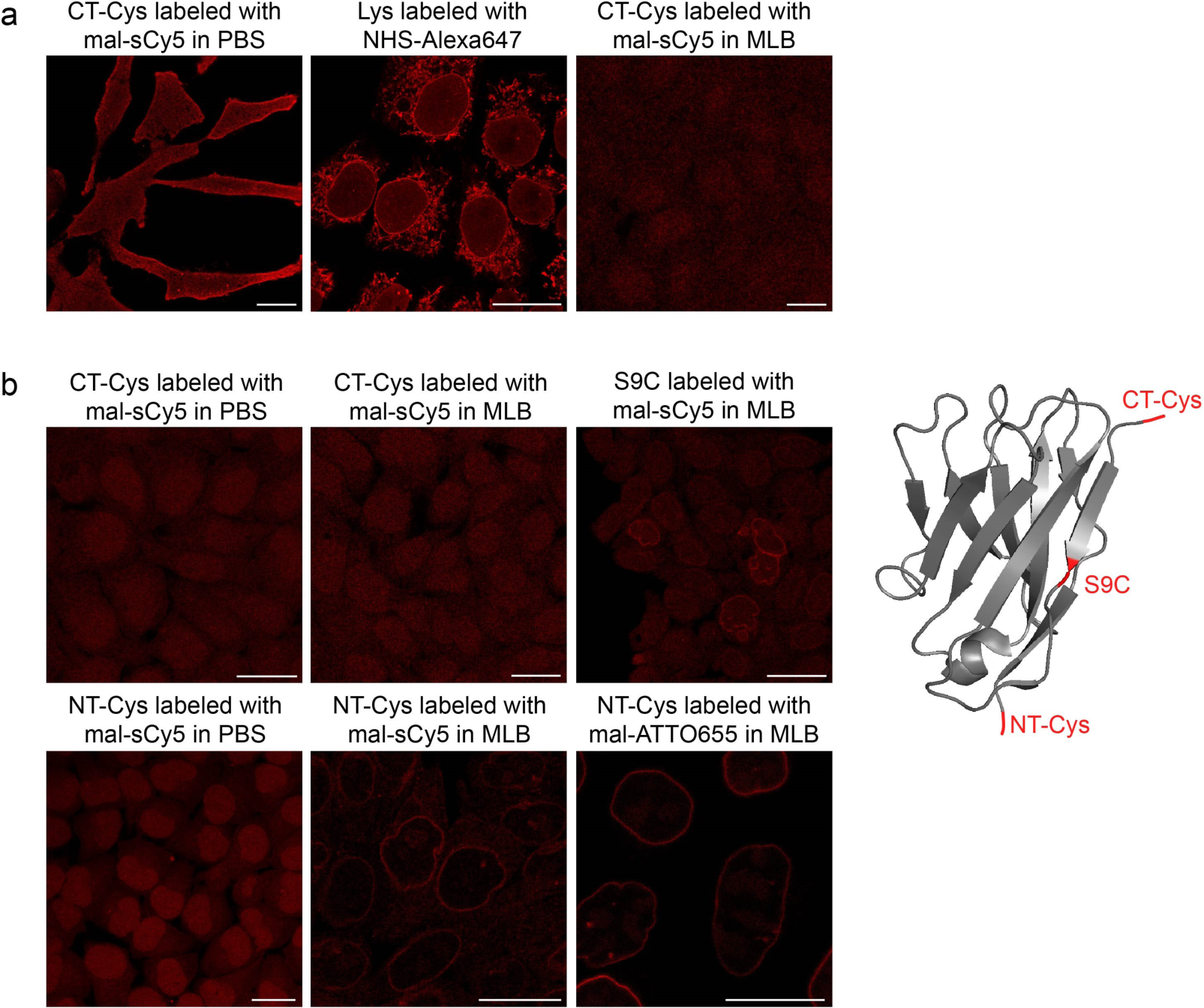
Labeling screen of differently modified α-Lamin fluorobodies. The α-Lamin nanobody was engineered with an extra single cysteine at three different positions, *e.g.* N-terminally (NT-Cys), C-terminally (CT-Cys), or at position 9 (S9C). After production in *E. coli* BL21/DE3 (**a**) or *E. coli* T7 SHuffle Express (**b**) and affinity purification, the engineered nanobodies were modified with maleimide sCy5 or ATTO655. Stochastic labeling via lysines exposed on the nanobody surface was conducted with NHS Alexa647. The specificity of these Fbs was evaluated by staining of HeLa Kyoto cells (500 nM) after fixation and permeabilization. Fb target specificity was analyzed by CLSM. For the CT-Cys labeled, S9C labeled and Lys labeled Fbs a severe lack in target specificity was observed. After production in *E. coli* T7 SHuffle Express and subsequent modification of the N-terminal cysteine in maleimide labeling buffer (MLB), specific tracing of endogenous lamin was achieved with greatly enhanced signal-to-background ratio. This Fb format was used for all further experiments. Scale bars: 20 µm.

**Figure 1 – Figure Supplement 5.**
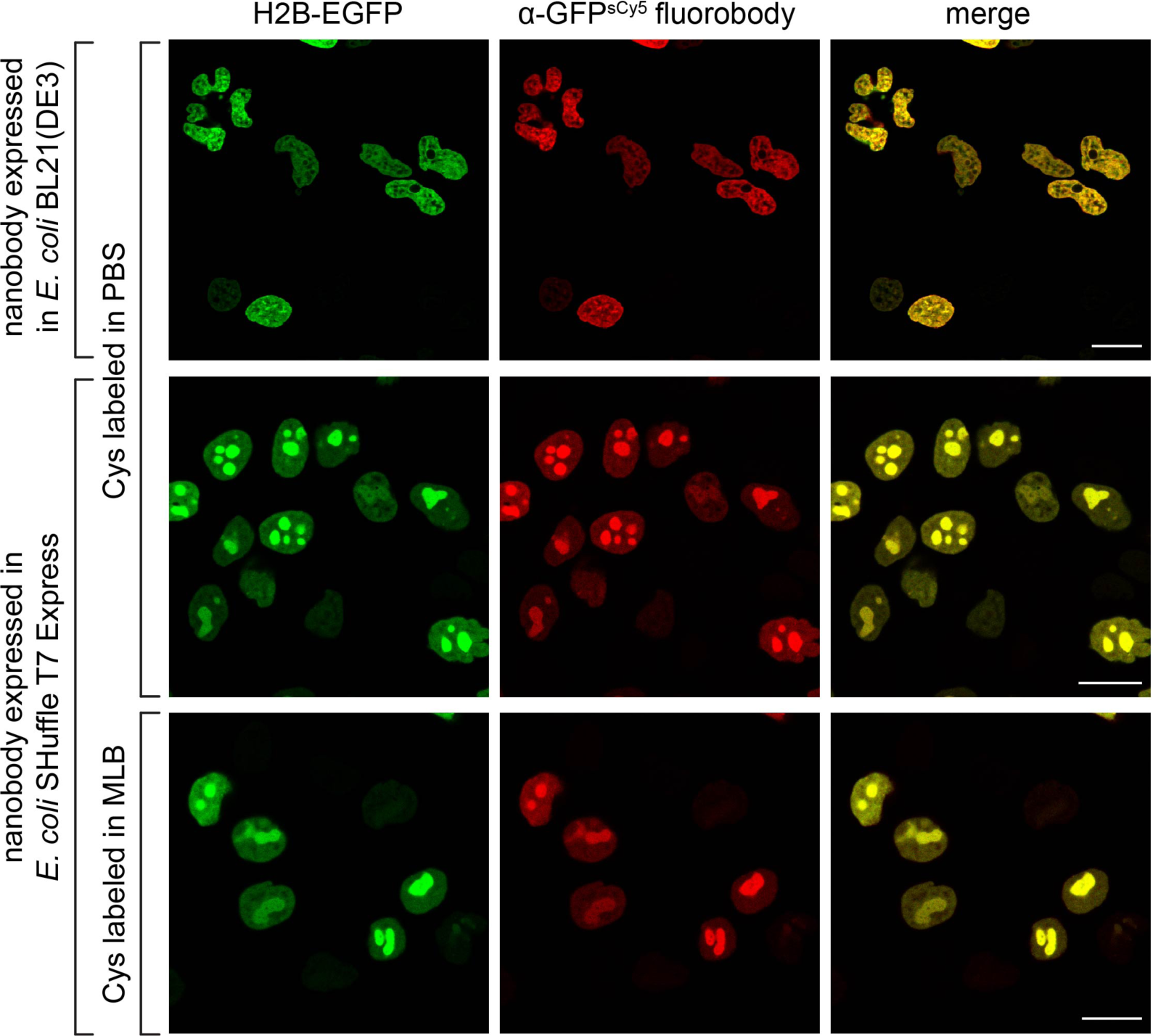
Ι Labeling screen of differently prepared α-GFP fluorobodies. The α-GFP^sCy5^ Fb with an N-terminal cysteine was produced in two different *E. coli* strains (BL21/DE3 or T7 SHuffle Express), and modified with maleimide sCy5. α-GFP binding screen was performed against H2B-EGFP (green) for all tested Fb preparation conditions. Fixed and permeabilized HeLa Kyoto cells were stained with α-GFP^sCy5^ Fb (100 nM, red). All combinations for Fb generation resulted in specific labeling of EGFP-tagged H2B (merge) by the engineered α-GFP with low background labeling. Analysis was performed by CLSM. Scale bars: 20 µm.

**Figure 2 – Figure Supplement 2.**
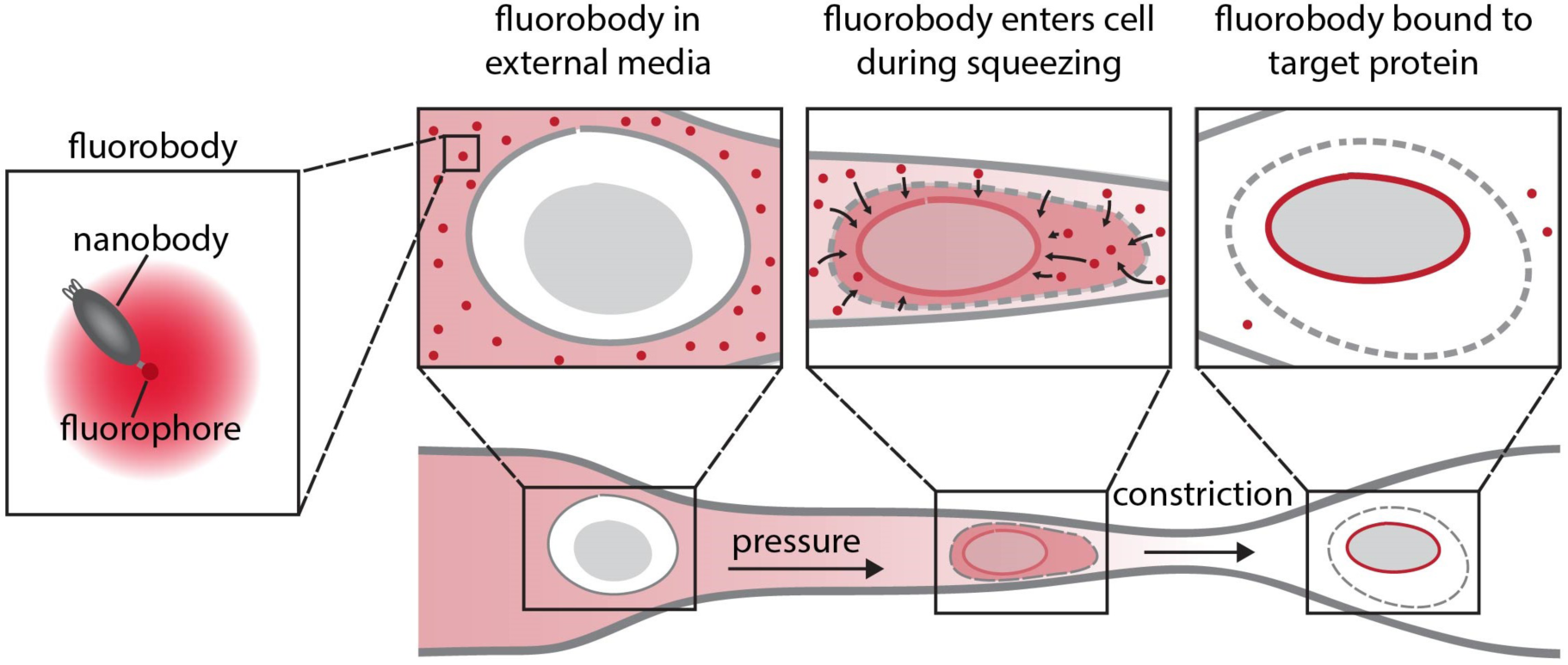
Fluorobody transfer into living cells by microfluidic cell squeezing. Cells are pressed through constrictions, which are ~ 30% smaller than cell diameter. Shear forces cause the formation of transient holes in the plasma membrane. By passive diffusion (middle), site-specifically labeled Fbs enter the cytosol and bind to their target (right, binding to nuclear envelope is exemplarily illustrated). The induced holes reseal within ~ 5 min.

**Figure 2 – Figure Supplement 2.**
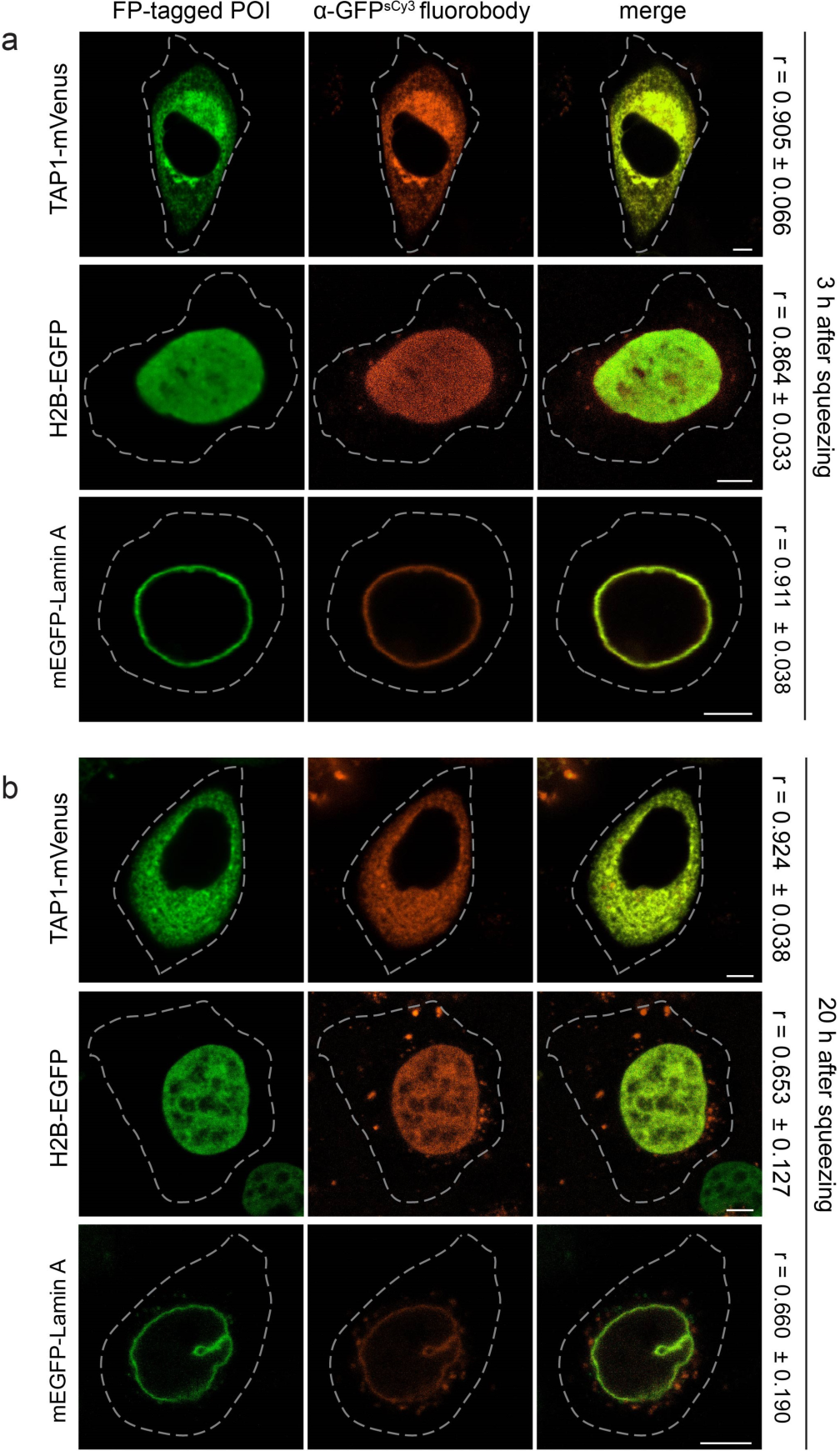
Live-cell labeling of FP-tagged targets via the α-GFP^sCy3^ fluorobody. α-GFP^sCy3^ Fb (200 nM) was transferred into HeLa Kyoto cells transiently transfected with TAP1-mVenus, H2B-EGFP, or mEGFP-Lamin A, respectively. Cells were imaged 3 h (**a**) and 20 h (**b**) after cell squeezing by CLSM. For all tested subcellular localizations of the FP-tagged POI, the molecular specificity of the Fb was persistent for a prolonged period of time (20 h). After 20 h, background originating as cytosolic punctae was detected. This observation is supported by a decreased Pearson´s coefficient (r, right), calculated from 8-10 individual cells. Dashed lines indicate the cell border. Scale bars: 5 µm.

**Figure 2 – Figure Supplement 3.**
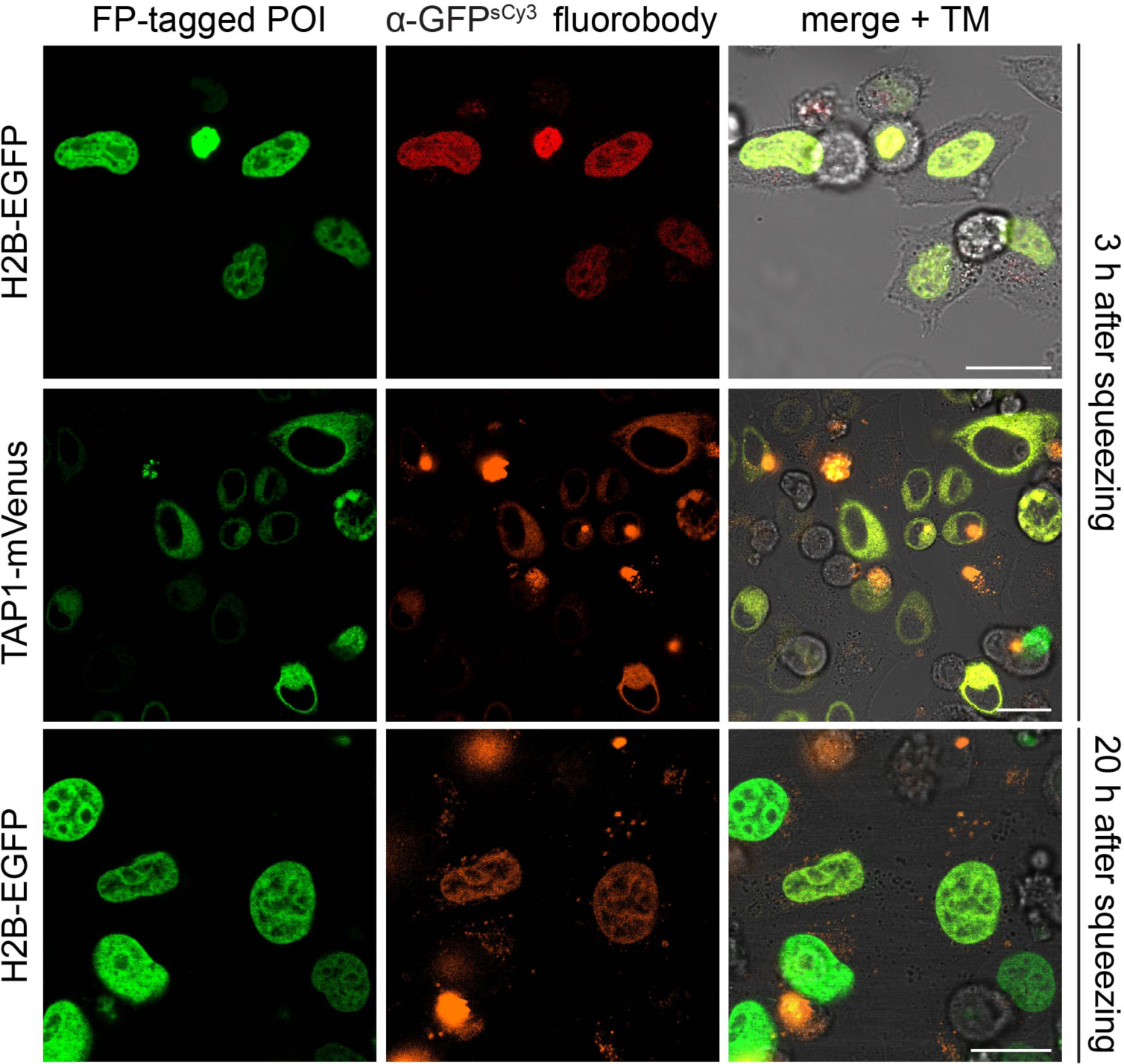
High fluorescence intensity correlation between target proteins and α-GFP^sCy3^ fluorobody in living cells. HeLa Kyoto cells expressing TAP1-mVenus or H2B-EGFP (green) were squeezed in the presence of α-GFP^sCy3^ Fb (200 nM, orange). Imaging via CLSM was performed 3 h and 20 h after squeezing. Low background and high co-localization between the Fb and POI was observed in multiple cells with varying expression level. Incipient punctual background was detectable after 20 h (bottom). Scale bars: 20 µm.

**Figure 2 – Figure Supplement 4.**
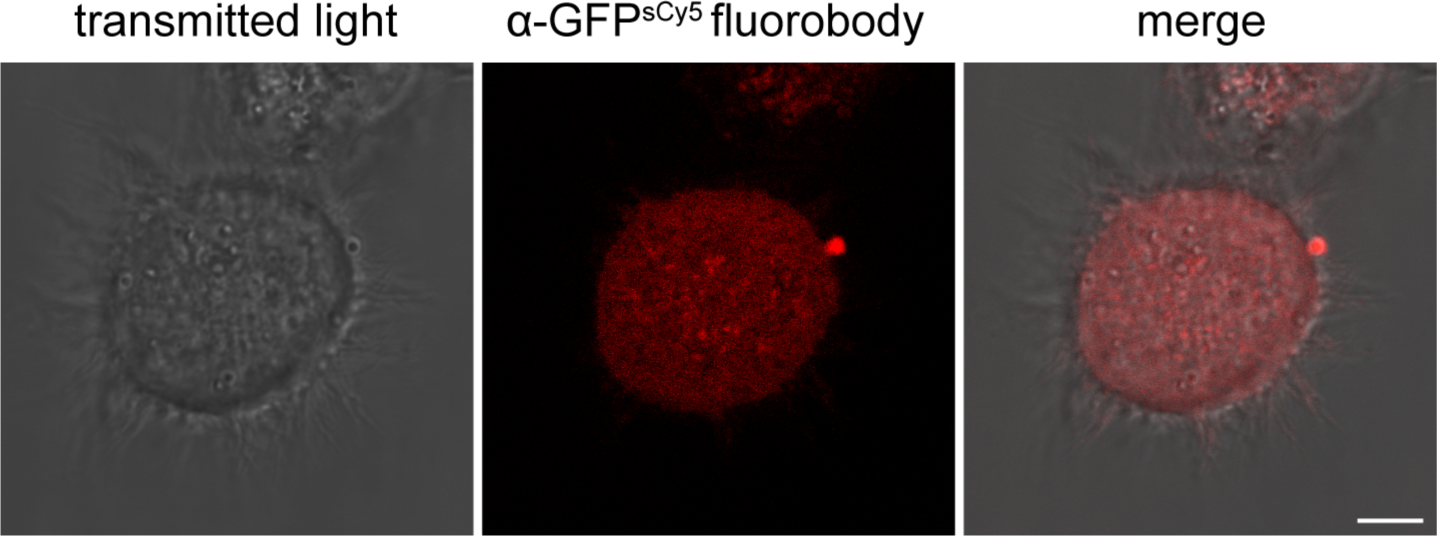
Cytosolic distribution of α-GFP^sCy5^ fluorobody in untransfected cells. α-GFP^sCy5^ Fb (100 nM) was delivered into HeLa Kyoto cells via cell squeezing. A cytosolic distribution of the Fb (red) was detected 1 h after squeezing. In absence of FP-tagged proteins, no distinct cellular localization of α-GFP^sCy5^ Fbs was observed, confirming the molecular specificity of the α-GFP^sCy5^ Fb for antigen tracking. Imaging was performed by CLSM. Scale bar: 5 µm.

**Figure 2 – Figure Supplement 5.**
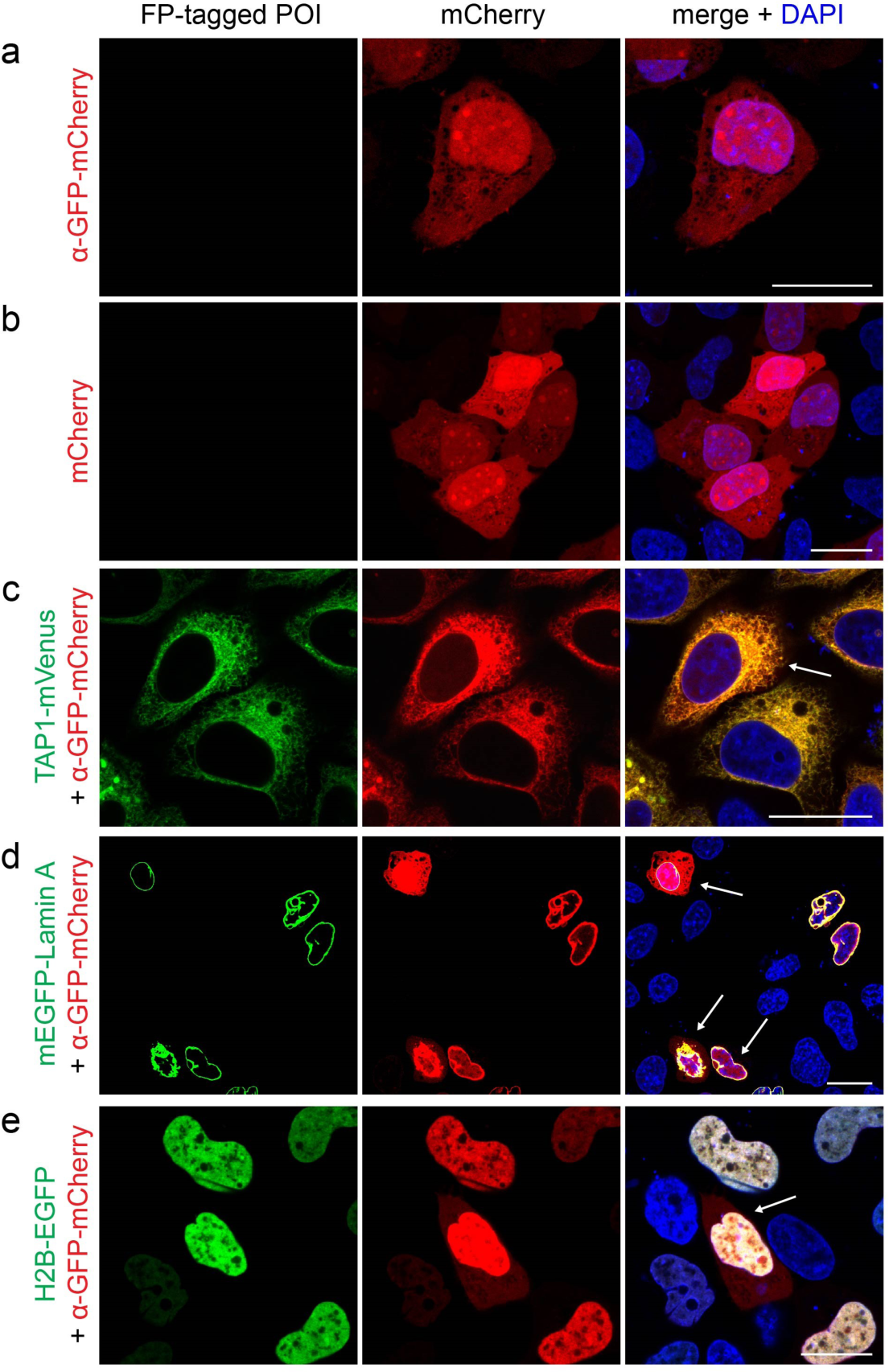
Ι Transient expression of α-GFP-mCherry chromobody in mammalian cells. (**a**) In the absence of the target protein, the α-GFP-mCherry nanobody was homogenously distributed within the cytosol and nucleus, similar to mCherry alone (**b**). (**c**-**e**) HeLa Kyoto cells were transiently co-transfected with an α-GFP-mCherry chromobody (red) and a FP-tagged POI (TAP1-mVenus, mEGFP-Lamin A or H2B-EGFP; green). Increased chromobody expression levels resulted in a high background of unbound nanobody, distributed in the cytosol and nucleus (indicated by arrows). Before CLSM imaging, HeLa Kyoto cells were chemically arrested with 4% formaldehyde and additionally stained with DAPI (blue). Scale bars: 20 µm.

**Figure 3 – Figure Supplement 1.**
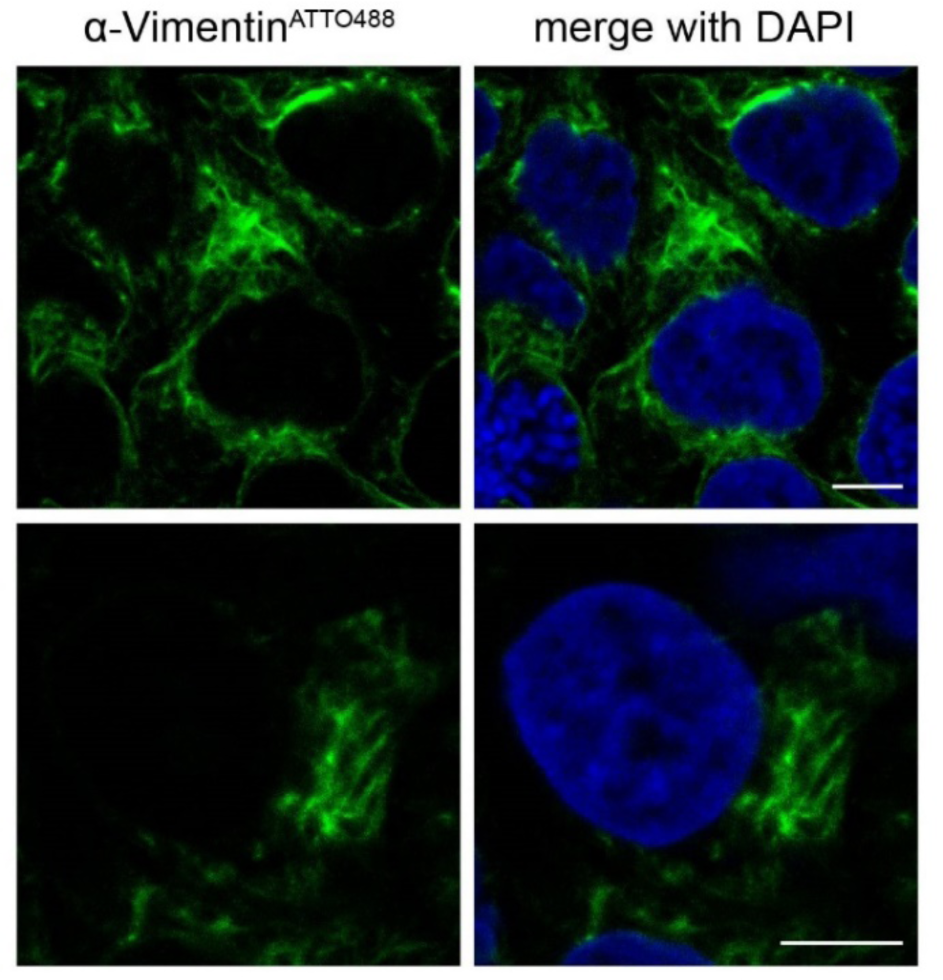
Vimentin staining in fixed cells. HeLa Kyoto cells were fixed, permeabilized and stained with α-Vimentin^ATTO488^ (20 µg/ml, green) and DAPI (blue). Confocal imaging showed specific labeling of endogenous filamentous vimentin. Scale bar: 5 µm.

**Figure 3 – Figure Supplement 2.**
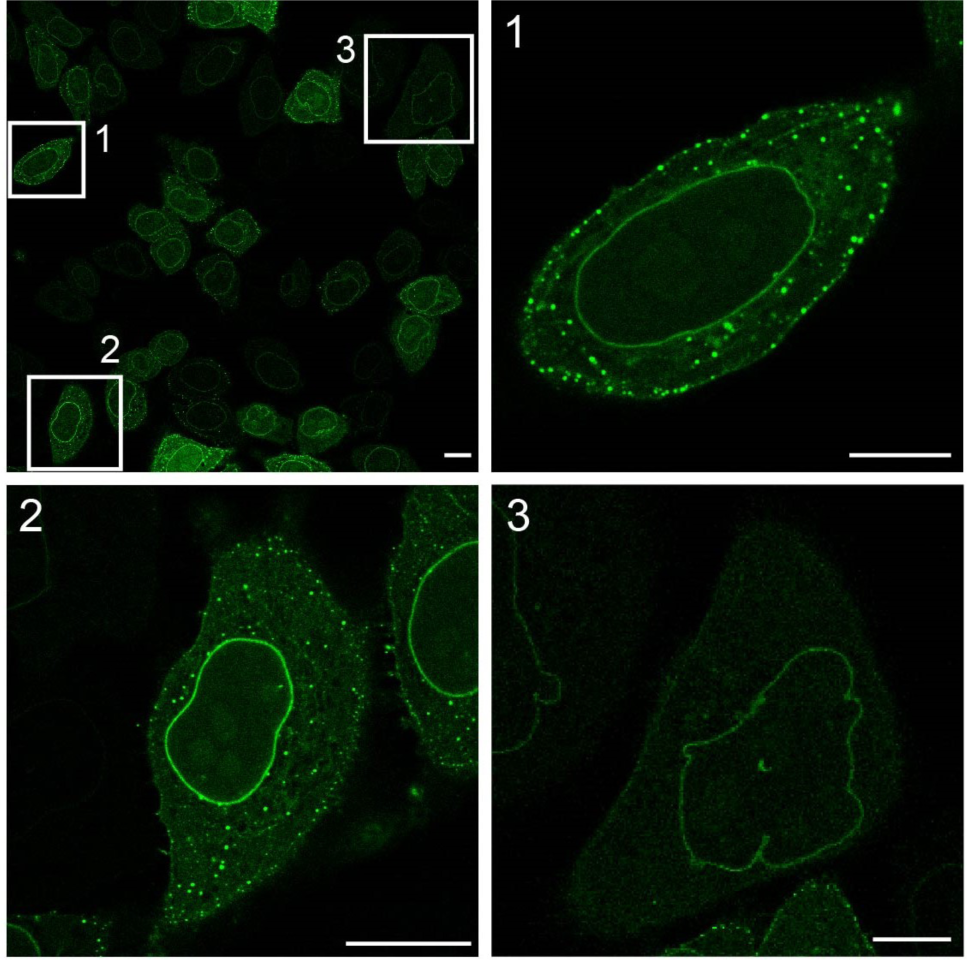
Transient expression of α-Lamin-EGFP chromobody in HeLa Kyoto cells entails background of free nanobody. Cells transiently transfected with α-Lamin-EGFP were fixed with 4% formaldehyde 24 h after transfection. Specific decoration of endogenous lamin at the nuclear envelope was visualized by CLSM. The variability of the chromobody expression level was accompanied by a high background, originating as cytosolic punctae. Images 1-3 are magnifications of individual cells as indicated by white boxes in the upper left image. Scale bars: 10 µm.

**Figure 3 – Figure Supplement 3.**
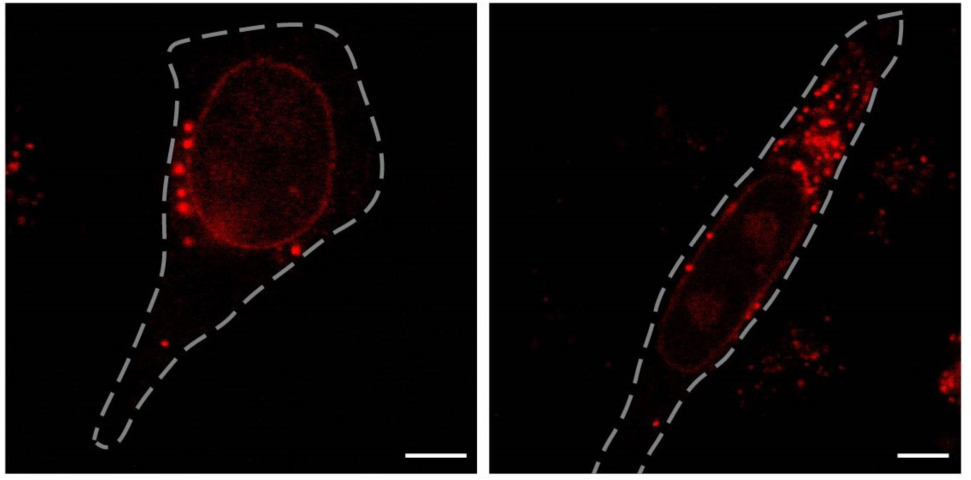
Persistence of endogenous lamin targeting by Fb in living cells. The α-Lamin^sCy5^ Fb (500 nM) was transferred into native HeLa Kyoto cells via cell squeezing. Specific decoration of the endogenous nuclear lamina by the Fb was still observed 20 h after squeezing by CLSM. An enriched punctual background was detected compared to 3 h post squeezing (see Fig. 3a). Dashed lines indicate the cell border. Scale bars: 5 µm.

**Figure 3 – Figure Supplement 4.**
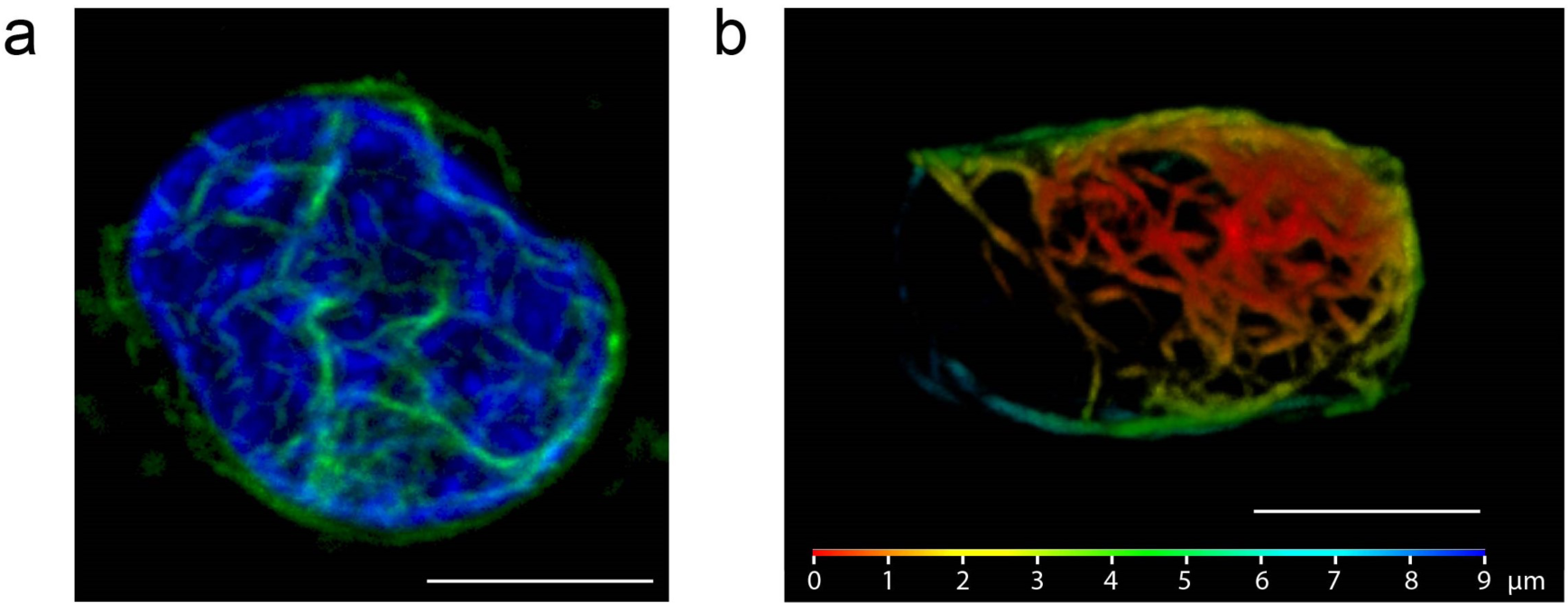
3D reconstruction of endogenous vimentin network. After intracellular transfer of α-Vimentin^ATTO488^ (50 µg/ml), z-stacks were recorded by confocal imaging using the Airy scan detector (SR mode, Zeiss). (**a**) Imaging was performed 3 h after squeezing with additional HOECHST staining (blue). The 3D reconstruction of the obtained z-stack showed the complex network of endogenous vimentin (green), organized around nucleus. (**b**) 3D topographic map of the endogenous vimentin meshwork. The series of z-stacks was recorded 20 h after cell squeezing and reconstructed. The rainbow-color code indicates the depth. Distance between stacks: ~ 200 nm, respectively. Scale bars: 5 µm.

**Figure 3 – Figure Supplement 5.**
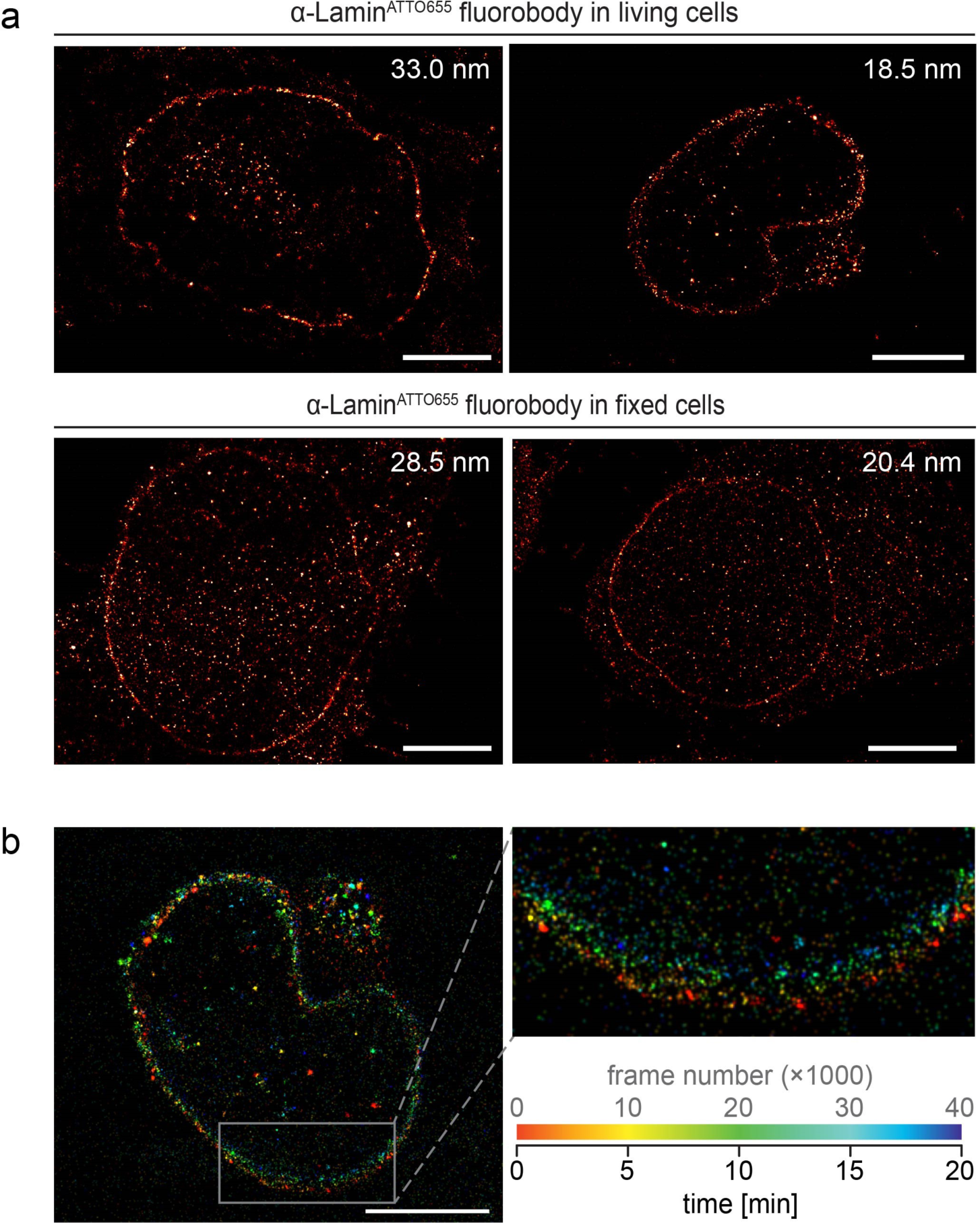
Super-resolution imaging of endogenous lamin in fixed and living cells. HeLa Kyoto cells were either squeezed with α-Lamin^ATTO655^ Fb (500 nM) for live-cell imaging or fixed, permeabilized and stained with α-Lamin^ATTO655^ Fb (100 nM). (**a**) *d*STORM analysis revealed a similar average localization precision in living (23.4 ± 8.3 nm, top) and fixed (23.2 ± 4.6 nm, bottom) cells. The localization precision determined for each image is given in the top right corner. Mean and standard deviation were calculated from three images. (**b**) Cell dynamics during live-cell super-resolution microscopy can impede imaging. Frame numbers are color-coded. Scale bar: 5 µm.

**Supplementary Video 1 Ι 3D reconstruction of the endogenous nuclear lamina.** After fixation and permeabilization, endogenous lamin of a HeLa Kyoto cell was visualized with an α-Lamin^ATTO655^ Fb (100 nM). 53 individual stacks were recorded by confocal imaging using the Airy scan detector (Zeiss, distance between stacks ~ 200 nm). 3D reconstruction showed a dense decoration of the complete nuclear lamina by the Fb, accompanied with a high signal-to-background ratio. Several intranuclear structures of endogenous lamin were observed.

